# Conserved exchange of paralog proteins during neuronal differentiation

**DOI:** 10.1101/2021.07.22.453347

**Authors:** Domenico Di Fraia, Mihaela Anitei, Marie-Therese Mackmull, Luca Parca, Laura Behrendt, Amparo Andres-Pons, Darren Gilmour, Manuela Helmer Citterich, Christoph Kaether, Martin Beck, Alessandro Ori

## Abstract

Gene duplication enables the emergence of new functions by lowering the general evolutionary pressure. Previous studies have highlighted the role of specific paralog genes during cell differentiation, e.g., in chromatin remodeling complexes. It remains unexplored whether similar mechanisms extend to other biological functions and whether the regulation of paralog genes is conserved across species. Here, we analyze the expression of paralogs across human tissues, during development and neuronal differentiation in fish, rodents and humans. While ~80% of paralog genes are co-regulated, a subset of paralogs shows divergent expression profiles, contributing to variability of protein complexes. We identify 78 substitutions of paralog pairs that occur during neuronal differentiation and are conserved across species. Among these, we highlight a substitution between the paralogs SEC23A and SEC23B subunits of the COPII complex. Altering the ratio between these two proteins via RNAi-mediated knockdown is sufficient to influence neuron differentiation. We propose that remodeling of the vesicular transport system via paralog substitutions is an evolutionary conserved mechanism enabling neuronal differentiation.

## Introduction

A major evolutionary event underlying the emergence of multicellular organisms is the specialization of functions between different cell types. An important role in defining the mechanisms that have led to this diversification is placed on the emergence of specific and definite gene expression programs that characterize distinct cell types (Arendt et al. 2016; Brunet and King 2017). Multicellular organisms are characterized by an increased genome complexity, in part driven by gene duplication events (Ohno 2013; Kaessmann 2010). Indeed paralog genes, namely genes that are the product of gene duplication events, are particularly enriched in the genomes of multicellular organisms (Lynch and Conery 2003). Even though in multicellular organisms the total paralog pool is generally larger, specific cell types express only a limited subset of paralogs, indicating the existence of mechanisms that restrict the expression of some paralogs genes in a given cell type (Padawer, Leighty, and Wang 2012). Most paralog genes share high sequence similarities and regulation of expression (Ibn-Salem, Muro, and Andrade-Navarro 2017). However, cases of divergent expression and regulation have been reported (Soria, McGary, and Rokas 2014;Makova 2003; Assis and Bachtrog 2015; Brohard-Julien et al. 2021), as exemplified by the distinct roles of Hox gene family members in modulating metazoan fronto-caudal development (Ferrier and Holland 2001). More recently, human specific gene duplications have been described to play a role in human brain development (Schmidt et al. 2019; Suzuki et al. 2018). Besides their modulation across cell types, an important role of paralogs is reflected by their ability to compensate for each other in maintaining the general homeostatic state of cells. Genome-wide CRISPR/Cas9-based screens have shown that paralog genes have a protective action on cell proliferation against the effect of gene loss-of-function in humans (Dandage and Landry 2019) and cancer cell lines (De Kegel and Ryan 2019; Thompson et al. 2021). All these observations highlight the functional impact that paralog genes have in modulating biological activity, development and cell differentiation.

From a molecular point of view, paralogs have been shown to modulate biological processes by influencing the assembly and activity of protein complexes. We have previously shown that specific compositions of protein complexes can be identified across cell types (Ori et al. 2016), and individuals (Romanov et al. 2019), and that the exchange of paralog complex members can contribute in specific cases to this variability. It has been also shown that the alternative incorporation of paralog proteins can antagonistically modulate the function of some protein complexes. For example, multiple specific paralog substitutions between subunits of the BAF chromatin remodelling complex lead to the assembly of functionally distinct complexes that can influence pluripotency and neuronal differentiation (Son and Crabtree 2014; Ho et al. 2009; Kaeser et al. 2008). Similarly, ribosomal paralog proteins promote ribosome modularity (Shi et al. 2017) and directly affect mRNA translation specificity (Gerst 2018; Slavov et al. 2015; Genuth and Barna 2018). Finally, co-expression analysis of protein complex members during human keratinocyte differentiation highlighted the existence of paralog subunits that compete for the same binding site in variable complexes (Toufighi et al. 2015). These studies indicate that paralog genes can contribute to the instalment of specific biological functions required, e.g., for cell differentiation, by influencing the activity of specific protein complexes. It remains currently unclear whether similar mechanisms extend to other molecular networks across the proteome and to which extent the regulation of paralog expression is conserved across cell types of different species.

In this study, using both newly generated and publicly available datasets, we systematically investigate how the expression of paralog genes contributes to transcriptome and proteome diversification across tissues, during development and neuronal differentiation. By integrating data from multiple organisms, we define a specific signature of paralog genes that emerges during neuronal differentiation and is conserved from fish to human.

## Results

### Co-expression of paralog genes during embryo development and across human tissues

In order to study the contribution of gene duplication to cell and tissue variability, we analyzed the expression profiles of paralog genes during zebrafish embryonic development and across healthy human tissues. We took advantage of two publicly available datasets describing a time-course transcriptome of zebrafish embryo development (White et al. 2017), and the steady state transcriptomes and proteomes of 29 healthy human tissues (Wang et al. 2019). We used correlation analysis of transcripts and proteins encoded by paralog genes to address their co-regulation during development and in fully differentiated tissues. According to Ensembl Compara (Yates et al. 2020) roughly 70% of the protein coding genes in the zebrafish and human genomes have paralogs, and similar proportions of paralogs are reflected in the datasets considered in this study (71% and 74% for zebrafish and human, respectively) (Fig1A). During zebrafish embryo development and across human tissues the majority of paralog genes pairs tend to be positively correlated (R > 0) (Table S1), however, a substantial proportion of them (33% and 36% for development and tissue, respectively) appears to be co-regulated in a negative manner (R ⇐0) (Fig1B, C). Comparable results were found also at the protein level across tissues, where the proportion of differently regulated paralog proteins appeared to be even higher (48%) (Supp.Fig1A). By calculating coefficient of variations for each protein and transcript, we also noticed that genes that possess paralogs in the genome tend to be more variably expressed during development (Supp.Fig1B) (two-sided Wilcoxon test p<2.2E-16), and across differentiated tissues at both transcriptome (Supp.Fig1C) (two-sided Wilcoxon test p<2.2E-16) and proteome level (Supp.Fig1D) (two-sided Wilcoxon test p<2.2E-16).

**Fig1.**
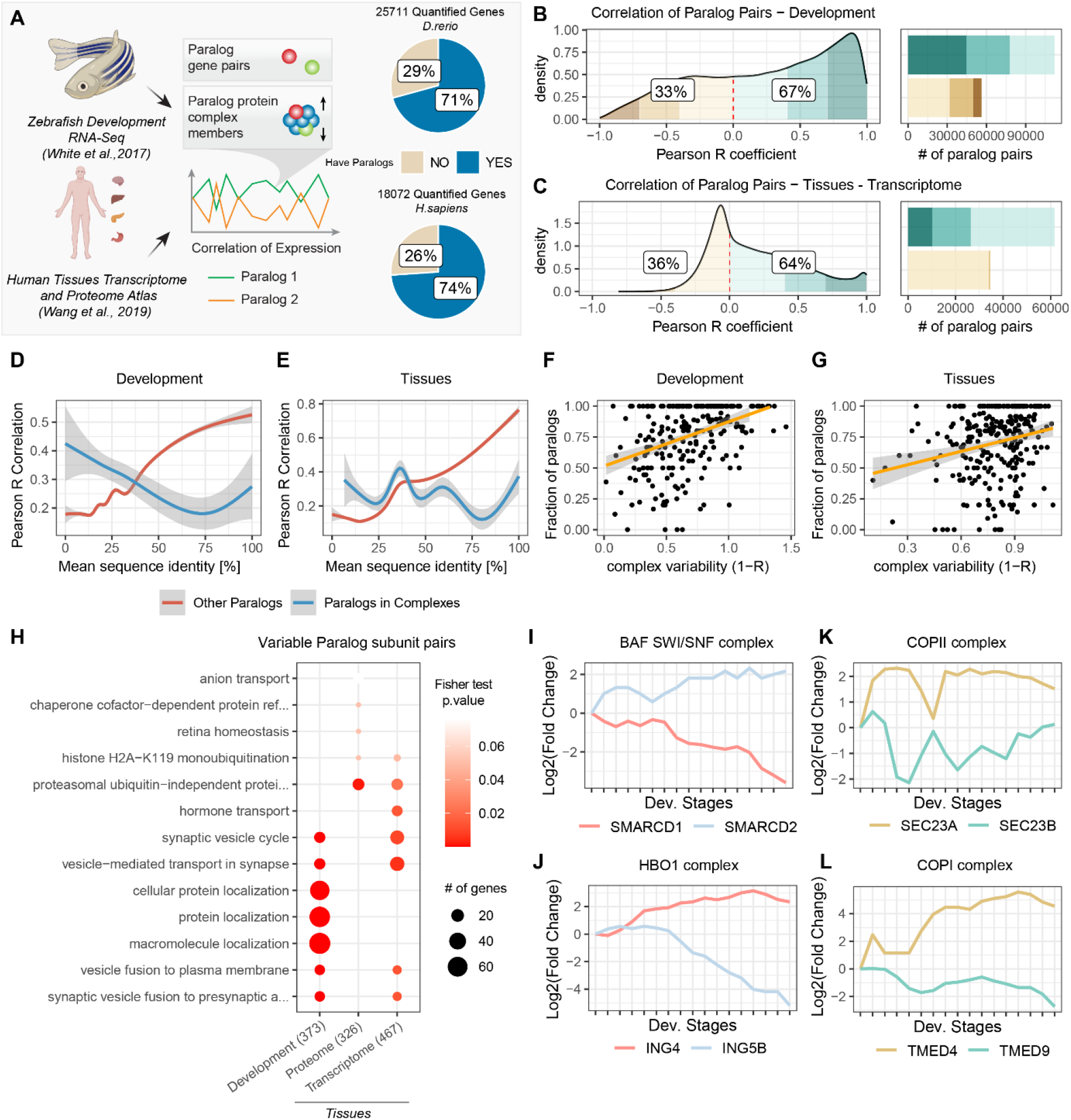
Expression of paralog genes during Zebrafish development and across human tissues. A - Transcriptome data during zebrafish embryo development (White et al. 2017) and transcriptome and proteome data from 29 healthy human tissues (Wang et al. 2019) were used to calculate Pearson correlation of expression during development and across tissues for paralog gene pairs. Pie plots indicate the proportion of quantified transcripts that possess at least one paralog in the zebrafish and human dataset, respectively. B/C - Density distribution of paralog gene pairs Pearson correlations during zebrafish embryo development (B) and across human tissues (C). Colored areas highlight different correlation intervals. Labels indicate the percentage of paralogs that are positively correlated (R=0) and negatively correlated (R⇐0). Barplot indicates the number of paralog pairs present in each category. Transcriptome data were used for both comparisons. D/E - Generalized Additive Model representing the relationship between mean paralog pairs reciprocal identity and transcript Pearson correlation in zebrafish development (D) and human tissues (E). Colored lines indicate paralog pairs that are members of the same protein complex (blue), and all other paralog pairs (red). F/G - Relationship between paralogs content (fraction of subunits that have paralogs in the genome) and complex variability. Complex variability is expressed as 1-R, where R is the median Pearson correlation of expression between all complex subunits. Transcriptome data were used for both embryo development (F) and tissues (G). H - GO term over representation analysis for the 25% most variable paralog subunit pairs against all other paralog subunit pairs. The top 5 most enriched terms from each dataset (development, tissues’s transcriptome, and tissues’s proteome) are shown. Numbers in parentheses on the x-axis indicate the number of unique variable paralog pairs considered for enrichment. I-L - Transcriptome profiles along embryo development for specific paralog pairs part of chromatin organization complexes, BAF SWI/SNF (I) and HBO1 (J), or vesicle-transport complexes, COPII (K) and COPI (L). Log2 fold changes calculated from TPMs relatively to the first time point are shown.

Since substitution of paralog members can contribute to the functional specialization of large protein complexes, such as chromatin remodeling complexes and ribosomes (Ori et al. 2016; Toufighi et al. 2015; Slavov et al. 2015; Romanov et al. 2019), we focused on the analysis of paralog expression in the context of protein complexes. We observed a characteristic behaviour of paralog pairs that assemble in the same protein complex. While paralogs co-expression was generally positively related to their sequence identity, i.e., highly similar paralogs tended to be co-regulated (R=0.16, Pearson correlation p=<2.2E-16 for development; R=0.33 Pearson correlation p<2.2E-16 for tissues), this was not the case for paralog pairs residing in the same protein complex, (R=−0.11, Pearson correlation p=3.69E-05 for development; R=−0.03, Pearson correlation p=0.19 for tissues, Fig1D, Fig1E, Table S1). This underlines the existence of a subset of paralog pairs that display no or negative co-expression, despite being members of the same protein complex and sharing a high sequence identity.

In order to estimate the contribution of these paralog genes to context-dependent protein complex formation, we investigated variations in the composition of macromolecular complexes during development and across tissues. We calculated the median correlation between all the possible pairs of genes belonging to the same protein complex and selected the upper and lower 25% percentiles of the resulting distribution to classify protein complexes as stable or variable, respectively (Supp.Fig2A, Table S2). During zebrafish development, we observed, as expected, positive correlations between protein complex members (Supp.Fig2B, p< 2.2E-16, two-sided Wilcoxon test). Highly correlated complexes include large house-keeping complexes, e.g., ribosomes and the proteasome, while functions carried out by more variable ones included molecular motors like the dynein-complex, vesicle associated proteins, e.g., SNARE, COPII/coat protein complex II, and chromosome and chromatin regulators, e.g., chromatin structure remodeling (RSC) complex (Supp.Fig2B, Table S2). The contribution of paralogs genes to the observed variability of protein complexes is highlighted by a general positive correlation between protein complex variability and paralog content, i.e., the fraction of complex members that have at least one paralog in the genome (R=0.40, p=2.1E-10) (Fig1F, Table S2). A similar pattern can be observed across human tissues at both transcriptome (R=0.23, p=7.9E-05, Fig1G, Table S2) proteome (R=0.27, p=1.3e-05) levels (Supp.Fig2C, Table S2). By calculating co-expression of single subunits (Supp.Fig2D), we consistently observed that complex members that possess at least one paralog tend to have a more variable expression compared to other members of the same complex (Supp.Fig2E, F, G, Table S2).

Interestingly, some of the most enriched Gene Ontology terms (GO) among anti-correlated paralog subunit pairs (bottom 25% of the distribution) were related to vesicle mediated transport and protein localization (Fig1H, Table S3), suggesting a potential divergent role of paralog proteins in establishing or modulating these biological functions. Our analysis recapitulated known anti-correlated expression for paralogs that are part of the BAF chromatin remodelling complex (homologous of the yeast SWI/SNF complex (Xue et al. 2000)) (Hansson et al. 2012; Ori et al. 2016; Ho et al. 2009) (Fig1I), but also specific expression profiles for members of the histone acetyl-transferase complex HBO1 (Fig1J), among others (Table S1). Similar expression patterns were observed also for paralogs belonging to complexes involved in the intracellular transport of macromolecules, such as the COPI and COPII complexes (Fig1K, Fig1L). Together these data suggest the existence of an evolutionary pressure for paralog subunits, especially involved in molecular trafficking and chromatin remodelling, to conserve sequence identity while diverging in expression across developmental stages and differentiated tissues.

### Conserved exchange of paralog proteins during neuronal differentiation

To investigate in more detail how the alternative usage of paralog genes contributes to cell variability, we focused on the well characterized process of neurogenesis that has been studied across different species by genome-wide approaches using both *in vivo* and *in vitro* model systems. We analysed neurogenesis datasets from zebrafish, mouse, rat and human (Fig2A, Table S4), based on the hypothesis that if some particular paralog substitutions are conserved across multiple organisms, they are more likely to functionally contribute to this process. We decided to use proteomics data to account for both transcriptional and post transcriptional mechanisms regulating paralog abundances.

**Fig2.**
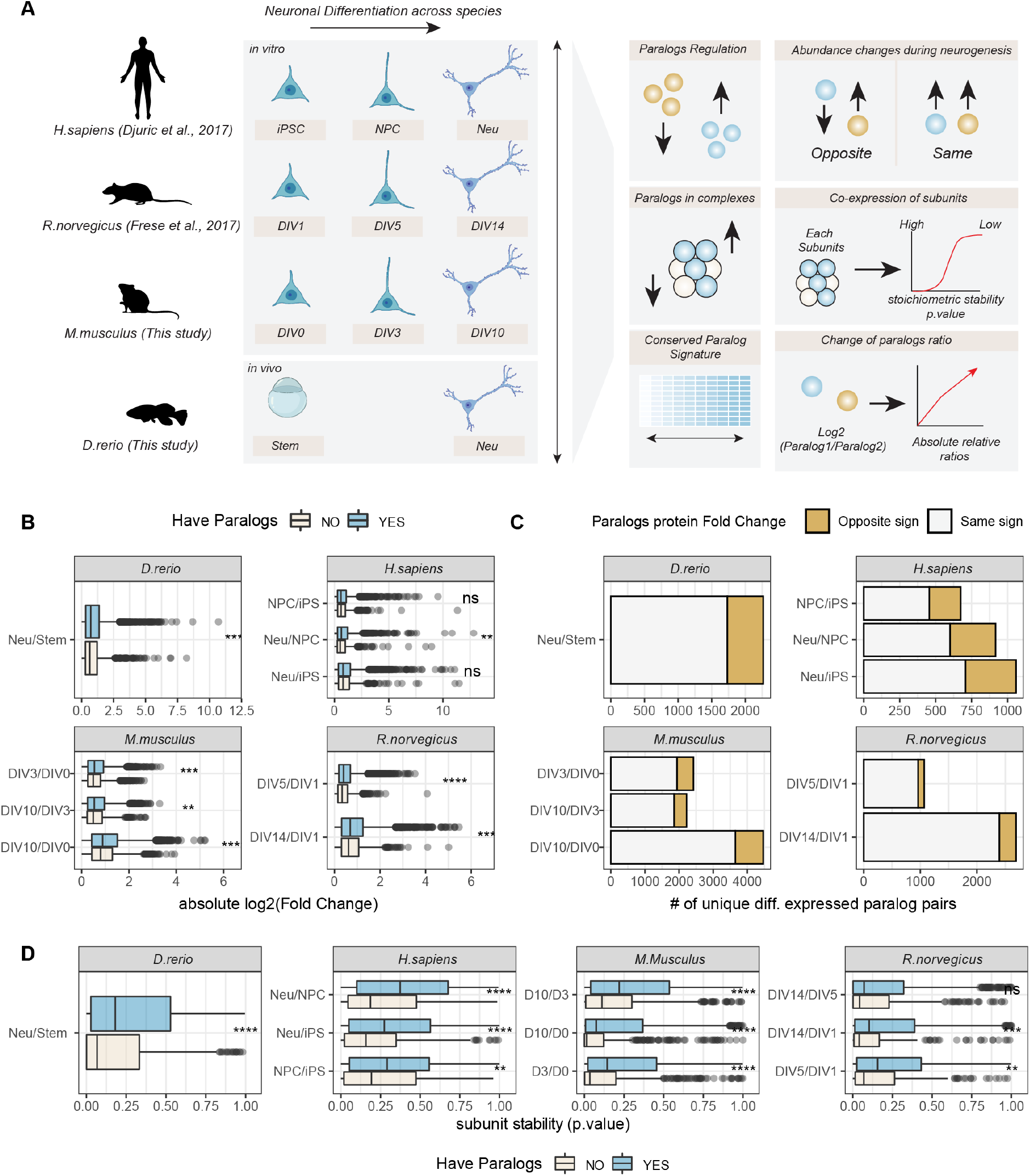
Changes of abundance of paralog proteins during neuronal differentiation. A - Overview of dataset used and data analysis workflow. DIV=differentiation *in vitro* day, iPSC=induced pluripotent stem cell, Neu=Neurons, NPC neuronal precursor cell, Stem=undifferentiated stem cell. B - Boxplots display absolute Log2 fold changes during neuronal differentiation for proteins that have (blue) or do not have (grey) at least one paralog. C - Barplots show the numbers of unique paralog pairs regulated in a concordant (grey) or opposite direction (orange) during neuronal differentiation. D - Boxplots compare the stability of protein complex subunits that have (blue) or do not have (grey) at least one paralog in the same protein complex. Low p values indicate subunits that are significantly co-expressed with the other members of the same protein complex and are therefore considered as “stable”. In B and D, asterisks indicate p values of the two-sided Wilcoxon test between the two compared groups: * p⇐0.05; ** p⇐0.01, *** p⇐0.001, **** p⇐0.0001, ns=not significant.

We generated a proteomic dataset using mouse primary neurons harvested after 0, 3 and 10 days of *in vitro* differentiation (DIV0, DIV3, DIV10). Shortly, cortical immature neurons were isolated from wild-type embryonic (E15.5) mouse brains and differentiated in glia-conditioned neurobasal medium. Neurons were collected at different time points and analysed by quantitative mass spectrometry (see Methods for details). We integrated this dataset with comparable data obtained from rat and human (Frese et al. 2017; Djuric et al. 2017). The rat dataset consisted of a time-course analysis of *in vitro* neurogenesis similar to the one performed in mouse, while the human data compared induced pluripotent stem cells (iPSC), iPSC-derived neural progenitor cells (NPCs) and cortical neurons (Neu). Finally, to directly compare the proteomes of embryonic stem cells and *in vivo* differentiated neurons, we took advantage of an established zebrafish line that enables the isolation of intact neurons using a fluorescent reporter. In this fish strain, the red-fluorescent-protein dsRed is expressed under the control of a neuronal-specific tubulin promoter from Xenopus (NBT-dsRed) (Peri and Nüsslein-Volhard 2008), allowing the selective isolation of neuronal cells by fluorescence-activated cell sorting (FACS). Undifferentiated cells were extracted from wild-type zebrafish 6 hours post fertilization (hpf), while NBT-dsRed zebrafish 1 day post fertilization (dpf) were used for the collection of differentiated neurons.

The quality of each dataset was evaluated using Principal Component Analysis (PCA) and GO enrichment analysis, confirming data reproducibility across replicates and the expected enrichment of terms related to neuronal development and cell differentiation (Supp.Fig3A, B, C, D). The neuronal differentiation data recapitulated the general patterns of paralog expression that we observed during development and across tissues: (i) proteins that have at least one paralog in the genome displayed larger fold changes (Fig2B); (ii) paralog pairs were generally co-regulated (Supp.Fig4A, Table S5); (iii) a subset of paralog pairs (~20%) displayed opposite regulation (Fig2C, Table S5). The latter set of paralogs was enriched for proteins related to chromatin remodeling, RNA splicing, RAS signalling, exocytosis and vesicle transport, as well as other processes related to development. Interestingly, while some enrichments were dataset-specific, we consistently observed an enrichment of GO terms related to DNA binding and transport across all datasets (Supp.Fig4B, Table S5). The neuronal differentiation datasets confirmed that paralogs contribute to protein complex variability, since in general, proteins that have at least one paralog display higher stoichiometric variability (Fig2D, Table S6), and, consequently, variable complexes were enriched in proteins with at least one paralog.

We then focused our analysis on paralog pairs that displayed divergent abundance changes during the neuronal differentiation process. In order to capture more subtle changes, we analyzed ratios between pairs of paralogs across conditions using absolute protein amounts estimated from mass spectrometry data. Briefly, log2 abundance ratios were calculated for all possible eggNOG (Huerta-Cepas et al. 2019) pairs across conditions, and significant changes in these ratios were statistically assessed using a linear model (see Methods) (Table S7). Differences in paralog ratios were sufficient to describe the general structure of the data, as highlighted by the separation of human, rodent and zebrafish dataset by PCA (Fig3A). By mapping every paralog pair to its relative eggNOG, we compared differences in paralog ratios across datasets. We were then able to assess which specific changes are conserved across the species considered. By applying a stringent cut-off (Log2 paralog ratio differences consistent in direction in all species and at least in 5 of the 7 conditions considered, and combined adjusted p⇐0.05, see Methods), we identified 78 paralog eggNOG pairs consistently affected during neuronal differentiation across all the species tested (Fig3B, Table S7). These conserved paralog pairs included multiple proteins involved in redox metabolism, RNA splicing, vesicles mediated trafficking and transport. Specifically, we found changes in ratios between the COPII complex subunits such as SEC23A and SEC23B (Fig3C), components of the retromer complex (VSP26B and VPS26A) (Supp.Fig5A), dynein subunits (DYNC1LI1 and DYNC1LI2) (Supp.Fig5B), and GTPase regulators of vesicle trafficking (RAB14 and RAB8A) (Supp.Fig5C). Taken together, these data highlight a potential role for paralogs proteins in mediating modularity of protein complexes during neuronal differentiation. Highly conserved substitutions between paralogs appear to predominantly affect paralog pairs that participate in the formation of transport complexes. This suggests that these substitutions might be required to adapt the transport system during neuronal differentiation and development in general.

**Fig3.**
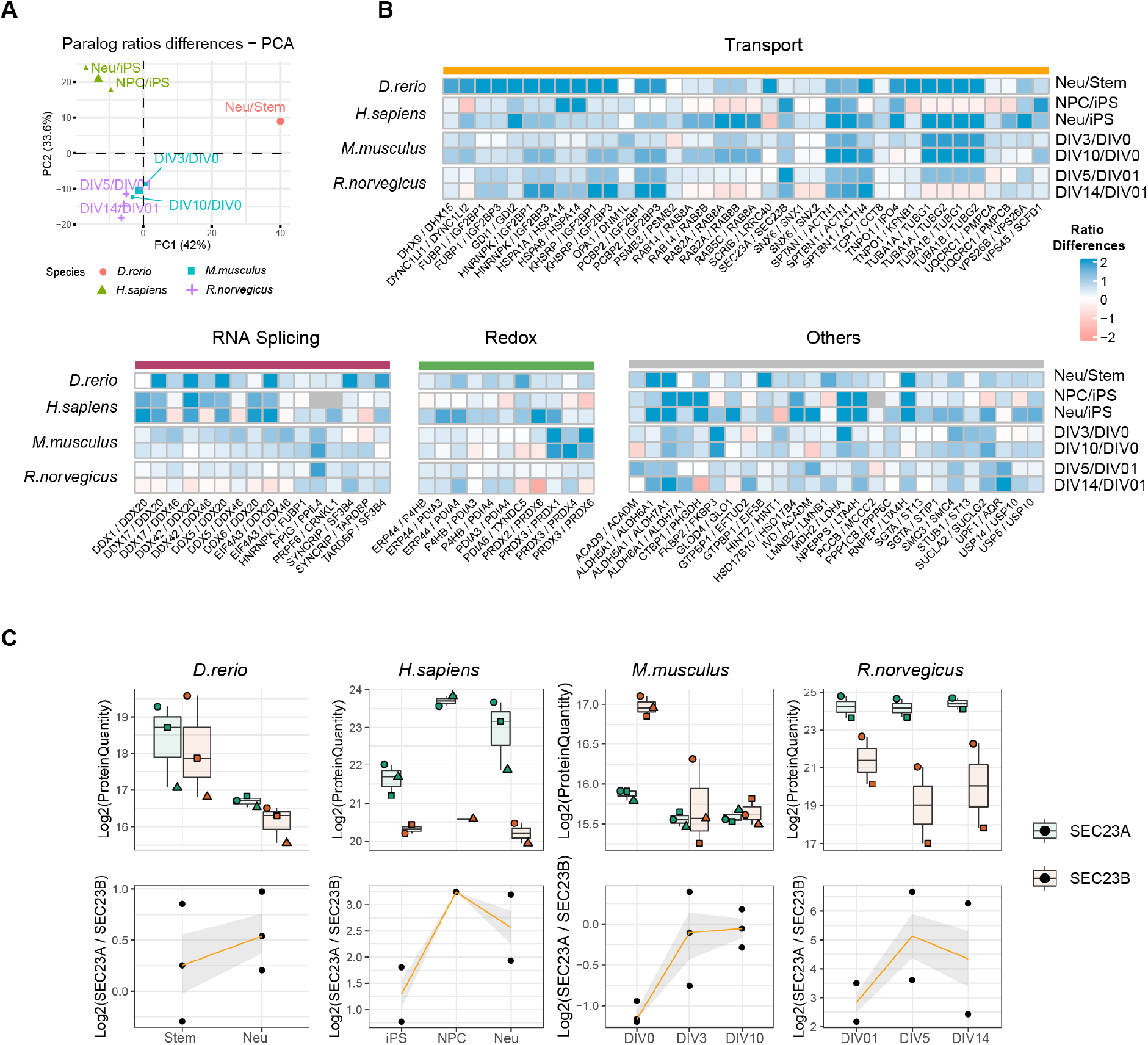
A conserved paralog signature during neuronal differentiation. A - Principal Component Analysis based on paralog ratio differences across conditions. Only paralog ratios quantified in all datasets are used for the analysis. The color code indicates the different species analysed, the small symbols indicate the different comparisons tested, and the large symbols indicate the centroid for each species. B - Heatmap shows conserved paralog substitutions during neuronal differentiation. Each column represents a specific eggNOG paralog pair mapped to the same human genes. Grey tiles indicate paralog pairs not quantified in the given condition. Paralog pairs are grouped according to their known biological function. C - Protein abundance profiles for SEC23A (green) and SEC23B (orange) across datasets. Boxplots indicate Log2 protein quantities, across different replicates, while line plots (bottom) indicate the ratios between the two paralogs. In the top panel, shapes indicate paired replicate experiments. In the bottom panel, orange lines indicate the mean paralog ratio across replicates, and the shaded area represents 50% confidence intervals.

### Altering the ratio between SEC23A and SEC23B affects neuronal differentiation

To experimentally test this hypothesis, we focused on the COPII subunits SEC23A and SEC23B. These are highly homologous paralogs that share a high level of protein sequence identity (>85%). The potentially divergent functions that these two particular paralogs may have are under debate (Zhu et al. 2015; Khoriaty et al. 2018), however they have never been studied in the context of neuronal differentiation. Using RNA interference (RNAi), we knocked-down either Sec23a or Sec23b in freshly isolated mouse neurons, and analyzed the respective proteome responses during *in vitro* neuronal differentiation (Fig.4A). First, we confirmed that RNAi significantly reduced the protein abundance of SEC23A relatively to a scrambled siRNA control (Fig4B, Log2 Fold Change siSec23a/siCtrl=−1.42, Qvalue=6.83 E-14, Table S8) and to a lesser extend for SEC23B (Log2 Fold Change siSec23b/siCtrl=−0.42, Qvalue=1.05 E-04, Table S8), globally altering the proportion between SEC23A and SEC23B in the differentiating cells. Interestingly, the knock-down of Sec23a induced a substantial compensatory increase of SEC23B (Log2 Fold Change siSec23a/siCtrl=1.06, Qvalue=6.06 E-10, Table S8), thereby maintaining the total amount of Sec23 (summed abundance of SEC23A and SEC23B) compared to siRNA control (Supp.Fig6A). A similar compensatory increase was true for the knock-down of Sec23b, although to a lesser extent (Fig4B).

**Fig4.**
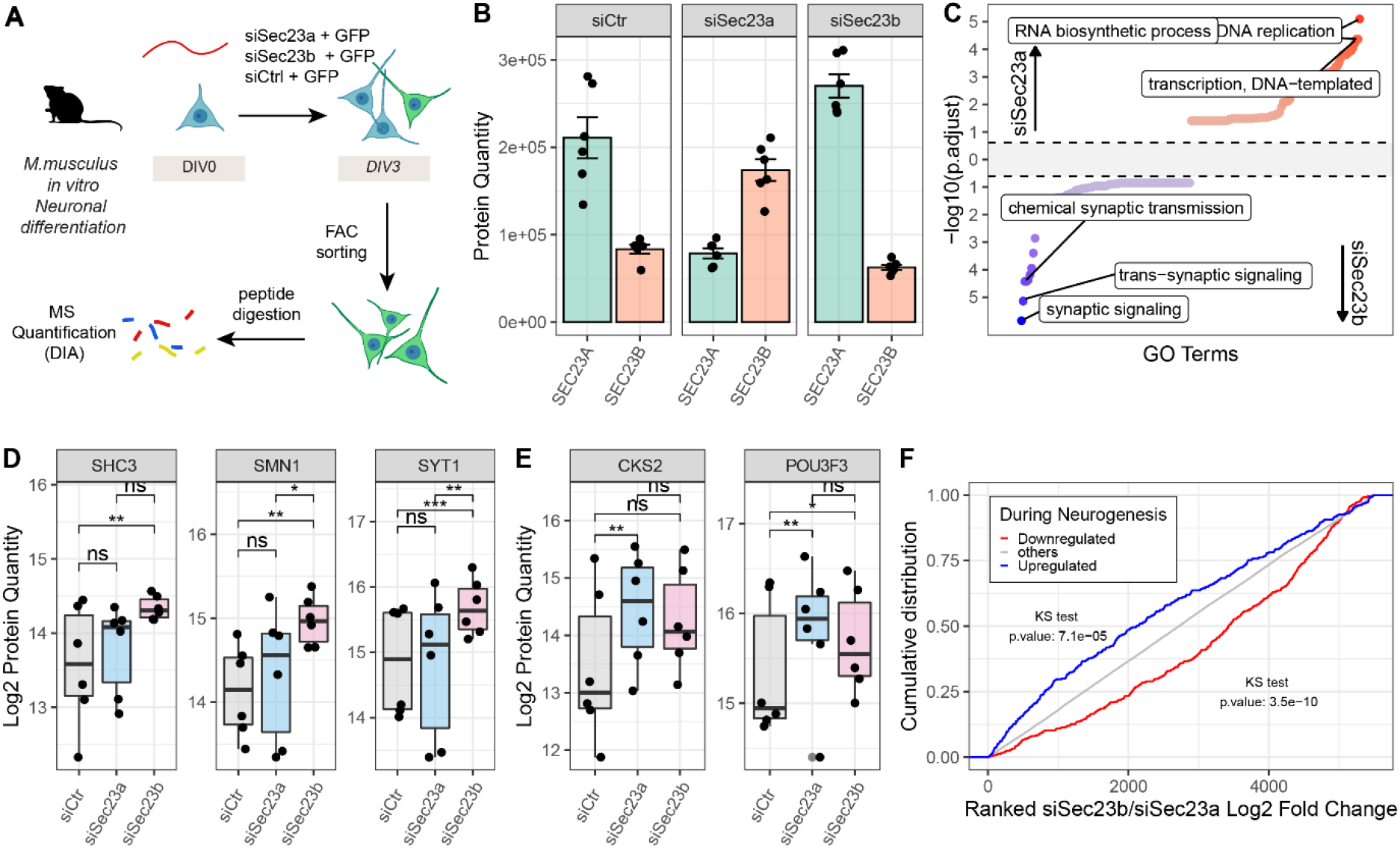
Altering the ratio between SEC23A and SEC23B affects neuronal differentiation *in vitro*. A - Mouse cortical neurons isolated from mouse embryos were transfected either with siCtr, siSec23a, or siSec23b, and a GFP expressing plasmid, and differentiated for 3 days. Transfected cells were isolated via FACS based on GFP expression and their proteomes analized by quantitative mass spectrometry (MS) using Data Independent Acquisition (DIA). B - Protein abundance of SEC23A (green) and SEC23B (orange) following different siRNA treatments, estimated from mass spectrometry data. n=6. C - Gene Set Enrichment Analysis (GSEA) for “Biological Process” category of differentially abundant proteins in siSec23b vs. siSec23a. The x axis represents the GO terms ranked by their −log10 adjusted p value, for the two conditions, while the y axis represents the −log10(adjusted p value) for each term. Top 100 GO terms enriched among proteins that are more abundant in the siSec23a or siSec23b condition are highlighted in red and blue, respectively. D, E - Quantification of selected proteins that were differentially affected by siSec23b and siSec23a. Asterisk indicates p values for the indicated comparison, as calculated from mass spectrometry data using Spectronaut (see Methods for details). * p⇐0.05; ** p⇐0.01, *** p⇐0.001, **** p⇐0.0001, ns=not significant. n=6. F - Cumulative distributions of ranked Log2 fold changes (siSec23b/siSec23a) for proteins that are upregulated (blue) (Log2 FoldChange DIV3/DIV0 >=1 and adjusted p⇐0.05), or downregulated (red) (Log2 FoldChange DIV3/DIV0 ⇐−1 and adjusted p⇐0.05) during mouse neuronal differentiation.

In order to understand the impact of an altered balance between the Sec23 paralogs on neuronal differentiation, we compared proteome responses of the different knock downs (KD). The changes in protein abundance caused by the Sec23a-KD or Sec23b-KD were globally correlated when compared to siRNA control (R=0.52, p< 2.2E-16). However, significant paralog-specific differences could be observed (Supp.Fig6B). GO enrichment analysis performed on the direct comparison of SEC23A-KD vs. Sec23b-KD showed that knock down of SEC23B increased the amount of proteins closely related to neuronal activity, i.e., synaptic signalling, whereas, knock down of Sec23a led to an increase of proteins related to DNA replication and RNA transcription (Fig4C, Table S8). Among these proteins, Sec23b knockdown increased the amount of SHC-transforming protein 3 (SHC3), a protein known to promote and regulate axon guidance (Pelicci et al. 2002), SMN1, a component of the Survival Motor Neuron complex, also linked to neurogenesis and neuronal differentiation (Liu et al. 2011; Lauria et al. 2020), and Synaptotagmin-1 (SYT1), a neuronal synaptic protein involved in neurotransmitter release (Coppola et al. 2001) (Fig4D). On the other hand, knock-down of Sec23a increased the expression of the Cyclin-dependent kinases regulatory subunit 2 (CKS2), a protein known to promote cell proliferation (Kang et al. 2009; Lin et al. 2016), and of the transcription factor POU3F3 that has been shown to be necessary for the earliest state of neurogenesis (Sugitani 2002; Dominguez, Ayoub, and Rakic 2013) (Fig4E). This pattern suggests that a higher proportion of SEC23A (as induced by the knockdown of SEC23B) promotes a more ‘neuronal’ state, while the opposite is true for the SEC23B paralog, which appears to promote a more undifferentiated and proliferative state. In order to investigate whether these responses were more global, we directly compared the effects of Sec23a-KD and Sec23b-KD to the early changes of the proteome that occur between DIV3 and DIV0 using our mouse neuronal differentiation dataset (Supp.Fig3C). The knockdown of Sec23a increased the levels of proteins that are downregulated during neuronal differentiation (Kolmogorov-Smirnov, KS test p=3.5E−10, Fig4F, Table S8). In contrast, the knockdown of Sec23b promoted an increase of proteins upregulated during neuronal differentiation (Kolmogorov-Smirnov, KS test p.value=7.1E−05, Fig4F, Table S8). This analysis confirms a functional divergence between these two paralogs, with SEC23A promoting, and SEC23B delaying mouse neuron differentiation *in vitro*.

## Discussion

In this study, we characterized the specific roles that paralog genes have in promoting transcriptome and proteome variability during development, neuronal differentiation and across different tissues. In accordance with the theory that paralog genes are main carriers of biological variability (Guschanski, Warnefors, and Kaessmann 2017; Ohno 2013), we found that genes that have paralogs are more often differentially expressed across tissues, during development and neuronal differentiation, indicating that they can be used as general descriptors of these specific biological states. New functional modules may then emerge in different cell types by gene duplication and subsequent functional divergence (Arendt et al. 2016; Ori et al. 2016). In agreement with this, we found that divergent expression is particularly pronounced for paralog gene pairs that participate in the formation of protein assemblies. More specifically, the disruption of the relationship between sequence identity and co-expression for this specific group of paralog genes could underline the existence of a specific evolutionary pressure to generate variable “modules” that interchange structurally to form distinct protein complexes with different functions. Consistently, stoichiometrically variable complexes are the ones with the highest paralog content, and they are often associated with functions related to membrane trafficking and chromatin organization. The observed modularity could be then comparable to what has been described for other cellular compartments, such as vertebrate synapses, where gene duplication of scaffold synaptic proteins has been related to the emergence of complex cognitive behaviours (Nithianantharajah et al. 2013).

Similar patterns of paralog regulation were conserved during neuronal differentiation in multiple vertebrates. By comparing data across species, we extracted a specific and conserved paralog exchange signature supporting the hypothesis of module divergence for membrane trafficking-related functions during neuronal differentiation. The relevance of paralog divergence in trafficking complexes has been also recently highlighted by the finding that two members of the COPI complex, COPG1 and COPG2, play distinct roles in modulating mouse neurogenesis (Jain Goyal et al. 2020). This specific substitution was also clearly identified in our mouse data, but not in all other datasets, suggesting that some of these functions could also be species specific. Moreover, we also addressed a similar exchange between the COPII complex members SEC23A and SEC23B that was highly conserved during neuronal differentiation from fish to humans. Previous studies on the functional divergence of these two paralogs reached contradicting conclusions, depending on the model system investigated. Some studies, carried out by substituting SEC23A in the SEC23B gene locus, have proposed a complete functional overlap of these two proteins (Khoriaty et al. 2018). Works by others have indicated separate roles regarding the ability to transport receptors (Scharaw et al. 2016) and cargo substrates (Zhu et al. 2015; Zeng et al. 2015). While these two paralogs are still highly redundant in function, we observed that they carry out different roles in respect to neuronal differentiation, with the SEC23A paralogs being needed to correctly progress during the neuronal differentiation process. Knockdown of either of the two paralogs induced opposite responses during *in vitro* neuronal differentiation, suggesting that a balanced paralog ratio is needed to correctly modulate this process.

More generally our study highlights the importance of paralog gene pairs in neuronal differentiation, as we have illustrated the possibility of promoting or antagonizing neuronal differentiation by targeting specific paralog genes. Similar mechanisms might be valid in other cell types or in different biological states, including pathological ones. Understanding which paralog genes define different cell identities could be exploited in the future for transdifferentiation purposes, e.g., for the generation of new models of neurodegenerative diseases (Mertens et al. 2018). In this case, we can speculate that specific paralog substitutions could help drive lineage transition between different somatic cells. However, broader comparisons between different cell types, integrating multiple data sources, single-cell analyses, and functional studies of specific paralogs are needed to better elucidate all these different possibilities.

## Acknowledgements

The authors gratefully acknowledge support from the FLI Core Facilities Proteomics, Flow Cytometry, Life Science Computing and Mouse, and the European Molecular Biology Laboratory (EMBL) Proteomics and FACS facilities. The authors thank Ivonne Heinze for processing samples for proteome analysis, Daniela Reichenbach and Christina Valkova for assistance with neuronal mouse culture, and Andrea Gruia for support with fish care. AO acknowledges funding from the German Research Foundation (Deutsche Forschungsgemeinschaft, DFG) via the Research Training Group ProMoAge (GRK 2155), the Else Kröner Fresenius Stiftung (award number: 2019_A79), the Fritz-Thyssen foundation (award number: 10.20.1.022MN) and the Chan Zuckerberg Initiative Neurodegeneration Challenge Network (NDCN). The FLI is a member of the Leibniz Association and is financially supported by the Federal Government of Germany and the State of Thuringia. CK acknowledges funding from the German Research Foundation (Deutsche Forschungsgemeinschaft, DFG, grant number KA 1751/8-1). MHC acknowledges funding from Fondazione AIRC per la Ricerca sul Cancro (AIRC, [IG 23539 to M.H.C.]).

## Author contributions

Conceptualization: DDF, LP, MB, AO. Data curation: DDF, LP. Experimental procedures: MA, MTM, LB, AAP, AO. Methodology: MA, MTM, AAP, AO. Project administration: AO, MB. Data analysis: DDF, LP. Supervision: AO, MB, CK, MHC, DG. Visualization: DDF. Writing – original draft: DDF, AO. Writing – review & editing: MA, MTM, LP, LB, MHC, CK, MB.

## Tables

**→ Table S1** - Zebrafish development data (White et al., 2017) and proteome and transcriptome data (Wang et al., 2019) across human tissues. This table also contains paralog pairs correlation values for those datasets.

**→ Table S2** - Protein complex and subunits co-expression during zebrafish development and across human tissues.

**→ Table S3** - GO Enrichment Analysis of variable subunit pairs during zebrafish development and across human tissues.

**→ Table S4** - Global proteomics data for neuronal differentiation in Mouse, Human, Rat and Zebrafish.

**→ Table S5** - Protein quantification data for paralog protein pairs during neuronal differentiation.

**→ Table S6** - Protein complex subunits co-expression during neuronal differentiation.

**→ Table S7** - Paralog pairs ratios during neuronal differentiation across species.

**→ Table S8** - Global proteomics data following SEC23A and SEC23B knockdowns during mouse neuronal differentiation.

## Materials and Methods

### Dataset and Resources

#### Ensembl Compara paralog genes resources

Paralogs annotation for *Homo sapiens* (GRch38.p13) *Danio rerio* (GRCz11)

*Mus musculus* (GRCm38.p6) *Rattus norvegicus* (Rnor_6.0), were downloaded from Ensembl (v102) via biomart (http://www.ensembl.org/biomart/martview/f04b3aa8b5c7f463e3edf9fa58d205a7).

Duplicated paralog pairs ( e.g, Paralog1 | Paralog2; Paralog2 | Paralog1) were removed from each dataset, so that only unique pairs (Paralog1 | Paralog2) were retained.

#### Protein Complexes Resources

Protein Complexes definition were taken from (Ori et al., 2016). Members of protein complexes were mapped by orthology in *Danio rerio* and *Rattus Norvegicus* using the bioconductor package ‘biomaRt’ (Durinck et al. 2009) using as reference the *Homo sapiens* protein complexes definitions.

#### Publicly Available Data used in this study

Zebrafish Embryo development data were obtained from White et al., 2017. (Supplementary file 3). Human Proteome and Transcriptome data across tissues were obtained from Wang et al., 2019 (Table EV2). Protein identification and LFQ intensity values (Log2) in cultured human iPSCs, NPCs and differentiated neurons, were obtained from Supplementary table S2 from (Djuric et al. 2017). Rat neuronal differentiation data published in Frese et al., 2017, were downloaded from PRIDE (http://proteomecentral.proteomexchange.org/cgi/GetDataset?ID=PXD005031) and analyzed again as described below.

### Isolation of embryonic stem cells and neurons from Zebrafish

Zebrafish (*Danio rerio*) strains were maintained following standard protocols (Westerfield, 2007) in the Gilmour lab at the EMBL, Heidelberg, Germany. Embryos were raised in E3 buffer (5 mM NaCl, 0.17 mM KCl, 0.33 mM CaCl2, 0.33 mM MgSO4) at 26 to 30 °C. All zebrafish experiments were conducted on embryos younger than 3 dpf. For isolation of undifferentiated cells a wild type strain (golden) and for neuronal cells the NBT-DsRed strain were used (Peri and Nüsslein-Volhard 2008).

#### Early embryos (6 hours post fertilization (hpf))

Wild type embryos were removed from their chorions using 1 ml of pronase (stock 30 mg/ml) in 40ml buffer E3 and incubated for 10-15 min with gentle shaking every 2 min in a small beaker. The supernatant was removed and the embryos were washed 4-5 times using buffer E3. The embryos were splitted into batches of around 250-300 per 1.5 ml tube. 1 ml of deyolking buffer (55 mM NaCl, 1.8 mM KCl, 1.25 mM NaHCO3) was added per tube and everything passed twice through a 200 μl pipet tip. The tubes were incubated at RT in a shaker at 1100 rpm for 5 min and afterwards spun at 300x *g* for 30 sec to remove the supernatant. The embryos were washed using 1 ml of wash buffer (110 mM NaCl, 3.5 mM KCl, 2.7 mM CaCl2 and 10 mM Tris/HCl pH 8.5), shaken at 1100 rpm at RT for 2 min and spun as above to remove the supernatant. The wash step was repeated twice. The deyolked and dissociated embryos were resuspended in 400 μl wash buffer and passed through a 40 μm cell strainer to remove undissociated cells. The merged cells were washed as above and resuspended in 110 μl PBS and counted using a hemocytometer.

#### Late embryos (24 hpf)

After 24 hpf, the NBT dsRed positive embryos were manually sorted. Up to the addition of the deyolking buffer all steps were the same as for early embryos. After the addition of 1 ml deyolking buffer per tube, the embryos were passed 10 times through a 1000 μl pipet tip, followed by washing twice with deyolking buffer and four times with washing buffer. For better cell dissociation the embryos were rinsed once with Accumax (Millipore) and then resuspended in 1ml Accumax and transferred to a 15 ml tube. The embryos were incubated at RT for 5 min at the lowest speed of the vortex mixer. The embryos were dissociated by pipetting for 2 min using a 1000 μl pipet tip, 2 min incubation on the vortex mixer and 1 min of additional pipetting. The cells were spun for 1 min at 300x *g* at RT and washed twice using 1 ml of PBS with 0.5 % BSA. 400 μl of PBS with 0.5 % BSA was used per tube to resuspend the cells afterwards passed through a 40 μm cell strainer and merged. DNAse I (Roche, 10 mg/ml in water) 170 U/ml and 10 mM MgCl2 was added. Cells expressing the DsRed fluorescent protein were FAC sorted with a MoFlo cell sorter (Beckman Coulter GmbH, Krefeld, Germany) to obtain a highly enriched fraction for neuronal cells.

### In vitro differentiation of mouse cortical neurons

#### Animal management practices

All mice were maintained in specific pathogen-free conditions, with food and water available ad libitum. The animal room had a constant temperature of 21°C ± 2, 55% ± 15 humidity, and controlled lighting (12 h light/dark cycle). The location for animal keeping was animal house TH4 at Leibniz Institute on Aging (Fritz Lipmann Institute), Jena, Germany. Breeding was license-free and performed under §11 TierSchG. Euthanasia and organ removal were performed under the internal §4 TierSchG licences O_CK_18-20 and O-CK_21-23. Euthanasia of mice was performed in a chamber with controlled CO2 fill rate according to “Directive 2010/63/EU annex IV of the European Parliament and the Council on the protection of animals used for specific purposes”.

#### Mouse neuronal cell culture

Cortical neurons were isolated from wild type murine embryonic brains (E15.5) of mixed background (FVB/NJ, C57BL/6, 129/Sv) and differentiated in glia-conditioned neurobasal medium. Briefly, meninges were removed, cortices were isolated, minced and dissociated in trypsin EDTA (Invitrogen), solution for 15 min at 37 °C. The supernatant was removed and the tissue was washed 3 times with trituration solution (10 mM HEPES, 1% penicillin/streptomycin, 10 mM L-glutamine, 1% BSA, 10% FBS, 0.008% DNase in HBSS) and homogenized in trituration solution using fire polished glass pipettes. For the mouse *in vitro* neuronal differentiation data, neurons were counted and pellets containing 1 million cells (DIV0) were prepared and frozen until further use. Additionally, 1 million cells were seeded on poly-L-lysine coated 6 cm plates containing 4 ml glia-conditioned plating medium (1% penicillin/streptomycin, 1 mM sodium pyruvate, 0.5% glucose, 10 mM HEPES 1x B27 supplement, 10% FBS, 10 mM L-glutamine in MEM). After 24 h the plating medium was substituted by glia-conditioned neurobasal medium (10 mM HEPES, 1x B27 supplement, 5 mM L-glutamine in NBM). Neurons were collected at DIV3 and DIV10. To this end, neurons were scraped off in cold PBS, obtained cell suspensions were transferred to a microcentrifuge tube and centrifuged for 5 min at 4 °C and 500 g. The obtained pellets were washed with PBS twice and frozen until further use.

For preparation of glia-conditioned mediums, a primary astroglia culture was established. For this purpose, brains were isolated from 15.5 days old embryos, the meninges were removed, the cerebral hemispheres were minced and afterwards dissociated in trypsin solution for 15 min at 37 °C. Finally, the tissue was homogenized by pipetting and cells were plated on a 10 cm dish containing glia medium (1% penicillin/streptomycin, 1 mM sodium pyruvate, 0.5% glucose, 10 mM HEPES, 20 mM L-glutamine, 10% FBS in MEM) and grown to confluence. For preconditioning of neurobasal medium or plating medium, the media were added to the glia feeder cultures and collected after 24 h.

#### SEC23A and SEC23B knockdown in mouse neuronal differentiation

For the Sec23 paralogs knockdowns, Cortical neurons were isolated from C57BL/6JRj mouse embryo (Janvier), as described above. Then, freshly isolated neurons (5 million cells per nucleofection reaction) were transfected using the 4D-Nucleofector™ X Unit and the P3 Primary Cell 4D Nucleofector X kit (Lonza, Switzerland), as indicated. Cells were transfected with 250 nM of siRNA and 1 μl of control pMax GFP (Nucleofector X kit, Lonza), using the CU-133 program. Immediately after transfection, cells were plated on poly-L-lysine (Sigma-Aldrich) - coated 10 cm plates containing 10 ml of glia-conditioned plating medium: 1% penicillin/streptomycin, 1 mM sodium pyruvate (Sigma- Aldrich), 0.5% glucose, 10 mM HEPES, 1x B27 supplement (Invitrogen), 10% FBS, 10 mM L-glutamine in MEM (Invitrogen 31095-052), and incubated at 37°C. After 1 day, the medium was replaced with glia-conditioned neurobasal medium: 10 mM HEPES, 1x B27 supplement, 5 mM L-glutamine in NBM (Invitrogen). After 3 days in culture, neurons were washed twice with PBS, detached using Trypsin EDTA (3-5 min, 37°C), collected in 5 ml of PBS with 2% FBS, and pelleted by centrifugation (450 g, 8 min, room temperature). Pellets were resuspended in 0.3 ml PBS with 2% FBS, and GFP positive cells were labeled with Sytox Blue Dead Cell Stain (viable staining) (Molecular Probes, ThermoFisher Scientific) and sorted directly in 200 μl of 2x lysis buffer (200 mM *HEPES pH 8.0*,100 mM *DTT,* 4% SDS) using a BD FACSAria Fusion with the Software BD FACSDiva 8.0.1 and 9.0.1 (BD Biosciences), using 488 nm laser and 530/30 filter for the GFP signal and laser 405 nm and 450/50 filter for the Sytox blue.

### Sample preparation for Mass Spectrometry

#### Sample preparation and dimethyl labeling for Zebrafish stem cells and neurons

Cells were lysed by addition of Rapigest (Waters) and urea to a final concentration of 0.2 % and 4 M, respectively, and sonicated for 3 x 30 sec to shear chromatin. Before protein digestion, samples were stored at −80 °C. Samples were quickly thawed and sonicated for 1 min. DTT was added to a final concentration of 10 mM and incubated for 30 min with mixing at 800 rpm to reduce cysteines. Then 15 mM of freshly prepared iodoacetamide (IAA) was added and samples were incubated for 30 min at room temperature in the dark to alkylate cysteines. Afterwards, 1:100 (w/w) LysC (Wako Chemicals GmbH) was added for 4 h at 37 °C with mixing at 800 rpm. Then urea concentration was diluted to 1.5 M with HPLC water and 1:50 (w/w) trypsin (Promega GmbH) was added for 12 h at 37 °C with mixing at 700 rpm. Afterwards the samples were acidified with 10 % TFA and the cleavage of Rapigest was allowed to proceed for 30 min at 37 °C. After spinning the sample for 5 min at 13,000x *g* at room temperature the supernatant was transferred to a new tube to proceed with peptide desalting.

For desalting and cleaning-up of the digested sample, C-18 spin columns (Sep-Pak C18 Classic Cartridge, Waters) were used. A vacuum manifold was used for all washing and elution steps. First the columns were equilibrated with 100 % methanol and then washed twice with 5 % (v/v) acetonitrile (ACN) and 0.1 % (v/v) formic acid (FA). The sample was loaded two times and then the column was washed 2 times with 5 % (v/v) ACN and 0.1 % (v/v) FA. The undifferentiated cell samples were labeled using a ‘light’ labeling reagent and the FACS sorted neuronal cells were labeled using an ‘intermediate’ labeling reagent inducing a mass shift of 28 or 32 Da respectively (Boersema et al. 2009). Formaldehyde and the D-isotopomer of formaldehyde react with primary amines of peptides (N-terminus and side chains of lysines) and generate a mass shift of 4 Da. The labeling reagents consisted of 4.5 ml 50 mM sodium phosphate buffer (mixture of 100 mM NaH2PO4 and 100 mM Na2HPO4), pH 7.5, 250 μl 600 mM NaBH3CN and 250 μl 4 % formaldehyde for light or 4 % deuterated formaldehyde for intermediate labeling reagent, per sample. After the labeling procedure, the column was washed 2 times with 5 % (v/v) ACN and 0.1 % (v/v) FA. For elution 50 % (v/v) ACN and 0.1 % (v/v) FA was used. Labelled peptides from undifferentiated cells and FACS sorted neurons were pooled, dried in a vacuum concentrator, and resuspended in 20 mM ammonium formate (pH 10.0), to be ready for high pH reverse-phase peptide fractionation. To dissolve the dried samples, they were vortexed, mixed for 5 min at maximum speed in a thermomixer and sonicated for 90 s. The samples were stored at −20 °C.

#### High pH reverse-phase peptide fractionation for dimethyl labelled samples

Offline high pH reverse-phase fractionation was performed using an Agilent 1200 Infinity HPLC System equipped with a quaternary pump, degasser, variable wavelength UV detector (set to 254 nm), peltier-cooled autosampler, and fraction collector (both set at 10°C). The column was a Gemini C18 column (3 μm, 110 Å, 100 x 1.0 mm, Phenomenex) with a Gemini C18, 4 x 2.0 mm SecurityGuard (Phenomenex) cartridge as a guard column. The solvent system consisted of 20 mM ammonium formate (pH 10.0) as mobile phase A and 100 % acetonitrile as mobile phase B. The separation was accomplished at a mobile phase flow rate of 0.1 ml/min using the following linear gradient: 99 % A for 2 min, from 99 % A to 37.5 % B in 61 min, to 85 % B in a further 1 min, and held at 85 % B for an additional 5 min, before returning to 99 % A and re-equilibration for 18 min. Thirty two fractions were collected along with the LC separation that were subsequently pooled into 10 fractions. Pooled fractions were dried in a speed-vac and resuspended in 5 % (v/v) ACN and 0.1 % (v/v) FA and then stored at −80 °C until LC-MS/MS analysis.

#### Sample preparation for in vitro differentiated mouse neurons

Frozen cell pellets of *in vitro* differentiated mouse neurons (~1 million cells per sample) were thawed and resuspended in 100 μl of 1x PBS. An equivalent amount of 2x Lysis Buffer (200 mM HEPES pH8.0, 100 mM DTT, 4% SDS) was added to the lysate, for a total volume of 200μl. For neurons treated with SEC23A/b or control siRNA, cells (between 40,000 and 180,000 cells) were sorted directly into 2x Lysis Buffer. Samples were then sonicated in a Bioruptor Plus (Diagenode, Seraing, Belgium) for 10 cycles with 1 min ON and 30 s OFF with high intensity at 20 °C. Samples were then boiled for 10min at 95°C, and a second sonication cycle was performed as described above. The lysates were centrifuged at 18,407x *g* for 1 min. Subsequently, samples were reduced using 10 mM DTT for 15min at 45°C, and alkylated using freshly made 15 mM IAA for 30 min at room temperature in the dark. Subsequently, proteins were precipitated using acetone and digested using LysC (Wako sequencing grade) and trypsin (Promega sequencing grade), as described in (Buczak et al. 2020). The digested proteins were then acidified with 10 % (v/v) trifluoroacetic acid. The eluates were dried down using a vacuum concentrator, and reconstituted samples in 5 % (v/v) acetonitrile, 0.1 % (v/v) formic acid. For Data Independent Acquisition (DIA) based analysis (siRNA treated neurons), samples were transferred directly to an MS vial, diluted to a concentration of 1 μg/μl, and spiked with iRT kit peptides (Biognosys, Zurich, Switzerland) prior to analysis by LC-MS/MS. For Tandem Mass Tags (TMT) based analysis (time course of *in vitro* differentiation), samples were further processed for TMT labelling as described below.

#### TMT labelling and high pH reverse-phase peptide fractionation

Following desalting, peptides were dried in a vacuum concentrator and buffered using 0.1M HEPES buffer pH 8.5 (1:1 ratio) for labelling, and then sonicated in a Bioruptor Plus for 5 cycles with 1 min ON and 30 s OFF with high intensity. 10-20 μg peptides were taken for each labelling reaction. TMT-10plex reagents (Thermo Scientific, Waltham, MA, USA) labeling was performed by addition of 1 μl of the TMT reagent. After 30 min of incubation at room temperature with shaking at 600 rpm in a thermomixer (Eppendorf, Hamburg, Germany), a second portion of TMT reagent (1μl) was added and incubated for another 30 min. After checking labelling efficiency, samples were pooled, desalted with Oasis® HLB μElution Plate and subjected to high pH fractionation prior to MS analysis. Offline high pH reverse phase fractionation was performed using a Waters XBridge C18 column (3.5 μm, 100 x 1.0 mm, Waters) with a Gemini C18, 4 x 2.0 mm SecurityGuard (Phenomenex) cartridge as a guard column on an Agilent 1260 Infinity HPLC, as described in (Buczak et al. 2020). Forty-eight fractions were collected along with the LC separation, which were subsequently pooled into 16 fractions. Pooled fractions were dried in a vacuum concentrator and then stored at −80°C until LC-MS/MS analysis.

### Mass Spectrometry data acquisition

#### Data Dependent Acquisition for dimethyl labelled samples (Zebrafish neurons and stem cells)

The 10 fractions obtained by high pH fractionation were analyzed using a nanoAcquity UPLC system (Waters GmbH) connected online to a LTQ-Orbitrap Velos Pro instrument (Thermo Fisher Scientific GmbH). Peptides were separated on a BEH300 C18 (75 μm x 250 mm, 1.7 μm) nanoAcquity UPLC column (Waters GmbH) using a stepwise 145 min gradient between 3 and 85% (v/v) ACN in 0.1% (v/v) FA. Data acquisition was performed using a TOP-20 strategy where survey MS scans (m/z range 375-1600) were acquired in the orbitrap (R=30,000 FWHM) and up to 20 of the most abundant ions per full scan were fragmented by collision-induced dissociation (normalized collision energy=35, activation Q=0.250) and analyzed in the LTQ. Ion target values were 1e6 (or 500 ms maximum fill time) for full scans and 1e5 (or 50 ms maximum fill time) for MS/MS scans. Charge states 1 and unknown were rejected. Dynamic exclusion was enabled with repeat count=1, exclusion duration=60 s, list size=500 and mass window ± 15 ppm

#### Data Dependent Acquisition for TMT labelled samples (mouse in vitro differentiation)

The 16 fractions obtained by high pH fractionation were resuspended in 10 μL reconstitution buffer (5% (v/v) acetonitrile, 0.1% (v/v) TFA in water) and 3 μL were injected. Peptides were separated using the nanoAcquity UPLC system (Waters) fitted with a trapping (nanoAcquity Symmetry C18, 5 μm, 180 μm× 20 mm) and an analytical column (nanoAcquity BEH C18, 2.5 μm, 75 μm× 250 mm). The outlet of the analytical column was coupled directly to an Orbitrap Fusion Lumos (Thermo Fisher Scientific) using the Proxeon nanospray source. Solvent A was water, 0.1% (v/v) formic acid, and solvent B was acetonitrile, 0.1% (v/v) formic acid. The samples were loaded with a constant flow of solvent A at 5 μL/min, onto the trapping column. Trapping time was 6 min. Peptides were eluted via the analytical column at a constant flow of 0.3 μL/min, at 40 °C. reconstitution buffer (5% (v/v) acetonitrile, 0.1% (v/v) TFA in water) and 3.5 μL were injected. Peptides were eluted using a linear gradient from 5 to 7% in 10 min, then from 7% B to 28% B in a further 105 min and to 45% B by 120 min. The peptides were introduced into the mass spectrometer via a Pico-Tip Emitter 360 μm OD ×20 μm ID; 10 μm tip (New Objective), and a spray voltage of 2.2 kV was applied. The capillary temperature was set at 300 °C. Full-scan MS spectra with mass range 375–1500 m/z were acquired in profile mode in the Orbitrap with resolution of 60,000 FWHM using the quad isolation. The RF on the ion funnel was set to 40%. The filling time was set at a maximum of 100 ms with an AGC target of 4 × 105 ions and 1 microscan. The peptide monoisotopic precursor selection was enabled along with relaxed restrictions if too few precursors were found. The most intense ions (instrument operated for a 3 s cycle time) from the full scan MS were selected for MS2, using quadrupole isolation and a window of 1 Da. HCD was performed with collision energy of 35%. A maximum fill time of 50 ms for each precursor ion was set. MS2 data were acquired with a fixed first mass of 120 m/z. The dynamic exclusion list was with a maximum retention period of 60 s and relative mass window of 10 ppm. For the MS3, the precursor selection window was set to the range 400–2000 m/z, with an exclude width of 18 m/z (high) and 5 m/z (low). The most intense fragments from the MS2 experiment were co-isolated (using Synchronus Precursor Selection=8) and fragmented using HCD (65%). MS3 spectra were acquired in the Orbitrap over the mass range 100–1000 m/z and resolution set to 30,000 FWMH. The maximum injection time was set to 105 ms, and the instrument was set not to injections for all available parallelizable time.

#### Data Independent Acquisition (SEC23A/b knockdowns)

Peptides were separated in trap/elute mode using the nanoAcquity MClass Ultra-High Performance Liquid Chromatography system (Waters, Waters Corporation, Milford, MA, USA) equipped with a trapping (nanoAcquity Symmetry C18, 5 μm, 180 μm × 20 mm) and an analytical column (nanoAcquity BEH C18, 1.7 μm, 75 μm × 250 mm). Solvent A was water and 0.1% formic acid, and solvent B was acetonitrile and 0.1% formic acid. 1 μl of the samples (∼1 μg on column) were loaded with a constant flow of solvent A at 5 μl/min onto the trapping column. Trapping time was 6 min. Peptides were eluted via the analytical column with a constant flow of 0.3 μl/min. During the elution, the percentage of solvent B increased in a nonlinear fashion from 0–40% in 120 min. Total run time was 145 min. including equilibration and conditioning. The LC was coupled to an Orbitrap Exploris 480 (Thermo Fisher Scientific, Bremen, Germany) using the Proxeon nanospray source. The peptides were introduced into the mass spectrometer via a Pico-Tip Emitter 360-μm outer diameter × 20-μm inner diameter, 10-μm tip (New Objective) heated at 300 °C, and a spray voltage of 2.2 kV was applied. The capillary temperature was set at 300°C. The radio frequency ion funnel was set to 30%. For DIA data acquisition, full scan mass spectrometry (MS) spectra with mass range 350–1650 m/z were acquired in profile mode in the Orbitrap with resolution of 120,000 FWHM. The default charge state was set to 3+. The filling time was set at a maximum of 60 ms with a limitation of 3 × 106 ions. DIA scans were acquired with 40 mass window segments of differing widths across the MS1 mass range. Higher collisional dissociation fragmentation (stepped normalized collision energy; 25, 27.5, and 30%) was applied and MS/MS spectra were acquired with a resolution of 30,000 FWHM with a fixed first mass of 200 m/z after accumulation of 3 × 106 ions or after filling time of 35 ms (whichever occurred first). Datas were acquired in profile mode. For data acquisition and processing of the raw data Xcalibur 4.3 (Thermo) and Tune version 2.0 were used.

### Mass Spectrometry data processing

#### Data processing for dimethyl-labelled samples (Zebrafish and Rat neuronal differentiation)

Software MaxQuant (version 1.5.3.28) was used to search the MS .raw data. For *D. rerio* the raw data were searched against the *D. rerio* UniProt database release: 2018_03, while for *R. norvegicus* the .raw files from (Frese et al. 2017), were downloaded from PRIDE repository PXD005031 and searched against the UniProt *R. norvegicus* database release 2019_08. Both datasets were searched appending a list of common contaminants. The data were searched with the following modifications: Carbamidomethyl (C) (fixed) and Oxidation (M) and Acetyl (Protein N-term; variable). For *D. rerio* 2 labels, Light L (DmethLys0 and DmethNterm0) and Heavy H (DmethLys4 and DmethNterm4) were selected representing the stem cell and neurons respectively. For the re-analysis of *R. norvegicus* data from Frese et al., 3 different labels were used: Light L (DmethLys0 and DmethNterm0), Medium M, (DmethLys4 and DmethNterm4) and Heavy H (DmethLys8 and DmethNterm8). For identification, match between runs was selected with a match time window of 2 minutes, and an alignment time window of 20 minutes. The mass error tolerance for the full scan MS spectra was set at 20 ppm and for the MS/MS spectra at 0.5 Da. A maximum of two missed cleavages was allowed. Identifications were filtered at 1% FDR at both peptide and protein levels using a target-decoy strategy (Elias and Gygi 2007). From each experiment, iBAQ values (Schwanhäusser et al. 2011) and ratios between labels were extracted from the ProteinGroups.txt table. Differential expression analysis was performed using the mean of the normalized ratios between labels. The R package fdrtool (Strimmer 2008) was used to calculate p values and q values for the different comparisons, on the Log2 transformed mean ratios.

#### Data processing for TMT10-plex data (mouse in vitro differentiation)

TMT-10plex data were processed using Proteome Discoverer v2.0 (Thermo Fisher Scientific, Waltham, MA, USA). raw files were searched against the fasta database (Uniprot *Mus musculus* database, reviewed entry only, release 2016_11) using Mascot v2.5.1 (Matrix Science) with the following settings: Enzyme was set to trypsin, with up to 1 missed cleavage. MS1 mass tolerance was set to 10ppm and MS2 to 0.5Da. Carbamidomethyl cysteine was set as a fixed modification while oxidation of methionine and acetylation (N-term) were set as variable. Other modifications included the TMT-10plex modification from the quantification method used. The quantification method was set for reporter ions quantification with HCD and MS3 (mass tolerance, 20ppm). False discovery rate for peptide-spectrum matches (PSMs) was set to 0.01 using Percolator 13 (Brosch et al. 2009). Reporter ion intensity values for the PSMs were exported and processed with procedures written in R (v.4.0.5) and R studio server (v.1.2.5042 and 1.4.1106), as described in (Heinze et al. 2018). Briefly, PSMs mapping to reverse or contaminant hits, or having a Mascot score below 15, or having reporter ion intensities below 1e3 in all the relevant TMT channels were discarded. TMT channels intensities from the retained PSMs were then log2 transformed, normalized and summarized into protein group quantities by taking the median value using MSnbase (Gatto and Lilley 2012). At least two unique peptides per protein were required for the identification and only those peptides with no missing values across all 10 channels were considered for quantification. Protein differential expression was evaluated using the limma package (Ritchie et al., 2015). Differences in protein abundances were statistically determined using the Student’s t test moderated by the empirical Bayes method. P values were adjusted for multiple testing using the Benjamini-Hochberg method (FDR, denoted as “adj. p”) (Benjamini and Hochberg, 1995).

#### Data processing for DIA samples (SEC23A/B knockdowns)

DIA libraries were created by searching the DIA runs using Spectronaut Pulsar (v13), Biognosys, Zurich, Switzerland). The data were searched against species specific protein databases (Uniprot *Mus musculus* release 2016_01) with a list of common contaminants appended. The data were searched with the following modifications: carbamidomethyl (C) as fixed modification, and oxidation (M), acetyl (protein N-term). A maximum of 2 missed cleavages was allowed. The library search was set to 1 % false discovery rate (FDR) at both protein and peptide levels. Libraries contained a total of 101,659 precursors, corresponding to 5708 and 6003 protein groups respectively. DIA data were then uploaded and searched against this spectral library using Spectronaut Professional (v.14.10) and default settings. Relative quantification was performed in Spectronaut for each pairwise comparison using the replicate samples from each condition using default settings, except: Data Filtering set to Qvalue sparse, and imputation to RunWise. Differential abundance testing was performed using a paired t-test between replicates. The data (candidate tables) and protein quantity data reports were then exported for further data analyses.

#### Data processing for human neuronal differentiation data

Protein identifications and LFQ intensity values (Log2) in cultured iPSCs, NPCs and differentiated neurons, were obtained from the original Supplementary table S2 published in (Djuric et al. 2017). Differential expression analysis between the different conditions was performed on the log2 LFQ intensity using the limma package (Ritchie et al. 2015).

### Data analysis

#### Analysis of paralog pairs during development and across tissues

For the Zebrafish development data (White et al., 2017), TPMs were used to calculate paralog pairs Pearson correlation coefficients. For the Human Tissue Atlas (Wang et al., 2019), Log2(FPKM) and Log2(IBAQ) were used to calculate correlation of paralog protein and transcript pairs. In all dataset only genes and proteins identified in at least 5 time-points / tissues were considered for correlation analysis. Coefficient of variations (σ / mean protein or transcript expression along time points / tissues) were also calculated for every gene in each datasets. Genes that have at least one paralog in the genome according to Ensembl Compara were labelled as ‘Have Paralogs’, and used for further analysis. From all the possible paralog pairs, 3 categories were created. The first one indicates all the possible paralog gene pairs, the second one indicates paralog pairs residing in the same protein complexes according to definitions from Ori et al., 2016, and the third one given by the exclusion between the two, indicating all other paralog pairs, namely paralog pairs that do not reside in the same complexes. For every paralog pair, the mean sequence identity was then calculated as the mean reciprocal identity retrieved from the Ensembl database. The relationship between sequence identity and co-expression between paralog pairs, was evaluated using Pearson R correlation coefficient, and visualized through a Generalized Additive Model.

#### Protein complex analysis during zebrafish development and across human tissues

For each datasets, proteins were annotated with the different protein complex definitions. Only protein complexes with at least 5 subunits present in each of the dataset were retained for analysis. For each of these complexes, all the possible pairwise correlations between subunits were considered, and from those the median value was used to calculate a median complex co-expression. We defined stable and variable complexes using the top and bottom 25% of the distribution respectively. (1-median Perason correlation) was also used to define then a measure of protein complex stoichiometric variability, as shown in Fig1F/G. The distribution of correlations was then compared with a distribution of randomly assembled complexes of same size and complex members obtained by randomly assigning proteins/transcripts to complexes. For each dataset, the fraction of paralog pairs present was considered as the number of subunits that have at least one paralog in the genome divided by the total size of each protein complex. Finally for each subunit, we calculated expression correlation values with the other members of the same complex, taking the median of this value as a measure of co-expression for that specific subunit. Top and bottom 25% of the obtained distribution were used to define stoichiometrically stable or variable subunits, respectively.

#### Paralog regulation during neuronal differentiation

For each datasets, differentially expressed proteins between different conditions (Log 2 Fold-Change > 0.58 and adjusted p value, or fdr tools p value < 0.05) were selected. Proteins were annotated as “Have Paralogs” if they had at least one paralog annotated in the genome. For each comparison, we then considered all possible paralog pairs present in the data and identified unique paralog pairs that displayed concerted regulation (same Log2 Fold Change sign for both paralogs) or opposite regulation (different Log2 Fold Change sign).

#### Subunits co-expression analysis for neuronal differentiation data

For calculating subunits stoichiometric variability we adapted a previously established pipeline (Gehring J, 2021). For each condition and datasets, only protein complexes that had at least 5 quantified subunits were considered. Then for each subunit in each complex, the median euclidean distance of fold change between that subunit and all other complex members was calculated. The distance obtained was compared with a distribution of distances for 2500 subunits from random complexes of equal size, obtained by randomly assigning proteins identified in the data to protein complexes. By comparing the two distributions we obtained a probability value for each subunit of observing lower distances with the complexes. Low p. values indicate high coexpression, denoted as stoichiometric stability, and vice versa.

#### eggNOG mapping

Fasta proteomes sequences used for MS protein quantification of the different dataset were annotated using emapper-2.1.4-2 (Cantalapiedra et al., 2021), based on eggNOG orthology data (Huerta-Cepas et al. 2019). Sequence searches were performed using the software MMseqs2 (Steinegger and Söding 2017). For each proteome eggNOG annotation was performed using default parameters.

#### Conserved exchange of paralog proteins

For each dataset, protein quantification values were used to calculate paralog ratios across conditions. The log2 paralog ratio between all possible quantified paralog pairs in each replicate was calculated for all the conditions tested. For each dataset, the significance of paralog ratio changes was assessed using the R package limma (Ritchie et al. 2015) considering replicates information. We considered only ratio changes relative to the first time point of each neuronal differentiation dataset. For comparison across species each paralog pair was mapped to its relative eggNOG. Only paralog pairs where both entries could be mapped to a valid eggNOG were retained. After eggNOG mapping, shared eggNOG pairs between species were used to assess if specific paralog substitution were shared across different organisms, and for each specific comparison we combined the p values using Fisher’s combined probability test from the metaRNASeq R package (https://cran.r-project.org/web/packages/metaRNASeq/index.html). Combined p values were corrected for multiple testing using the Benjamin-Hochberg correction (Benjamini and Hochberg 1995). Since in some cases multiple proteins can map to the same eggNOG, for each pair and condition the mean value was considered for both ratio differences and p values. From this analysis, we considered as “conserved” only paralog gene pairs identified in all species and whose log2 ratio changes were consistent in sign in at least 5 of the 7 neuronal differentiation comparisons, with combined adjusted p⇐0.05.

#### GO enrichment analysis

Over representation analysis of GO terms was performed with the R package topGO (https://bioconductor.org/packages/release/bioc/html/topGO.html). Fisher test was used in order to estimate the expected proportion for different terms and obtain a p value indicating the enrichment score for each specific GO term. Gene set enrichment analysis (GSEA) was performed with the topGO R package using a Kolmogorov-Smirnov test on the cumulative ranked distributions. For both enrichments p values were adjusted using Hommel’s correction, GOTerms were considered significant if their adjusted p values were below the value of 0.05. The R package rrvgo (https://ssayols.github.io/rrvgo/) was used in order to summarize and reduce redundancy of the enriched GO terms using default settings.

#### Figure generation

Data visualization was performed with R (v.4.0.5) and R studio server (Version 1.4.1106) using the ggplot2 package (Wickham 2009). Figure panels 1A, 2A, 4A were created with https://BioRender.com.

---

**Supp. Fig1.**
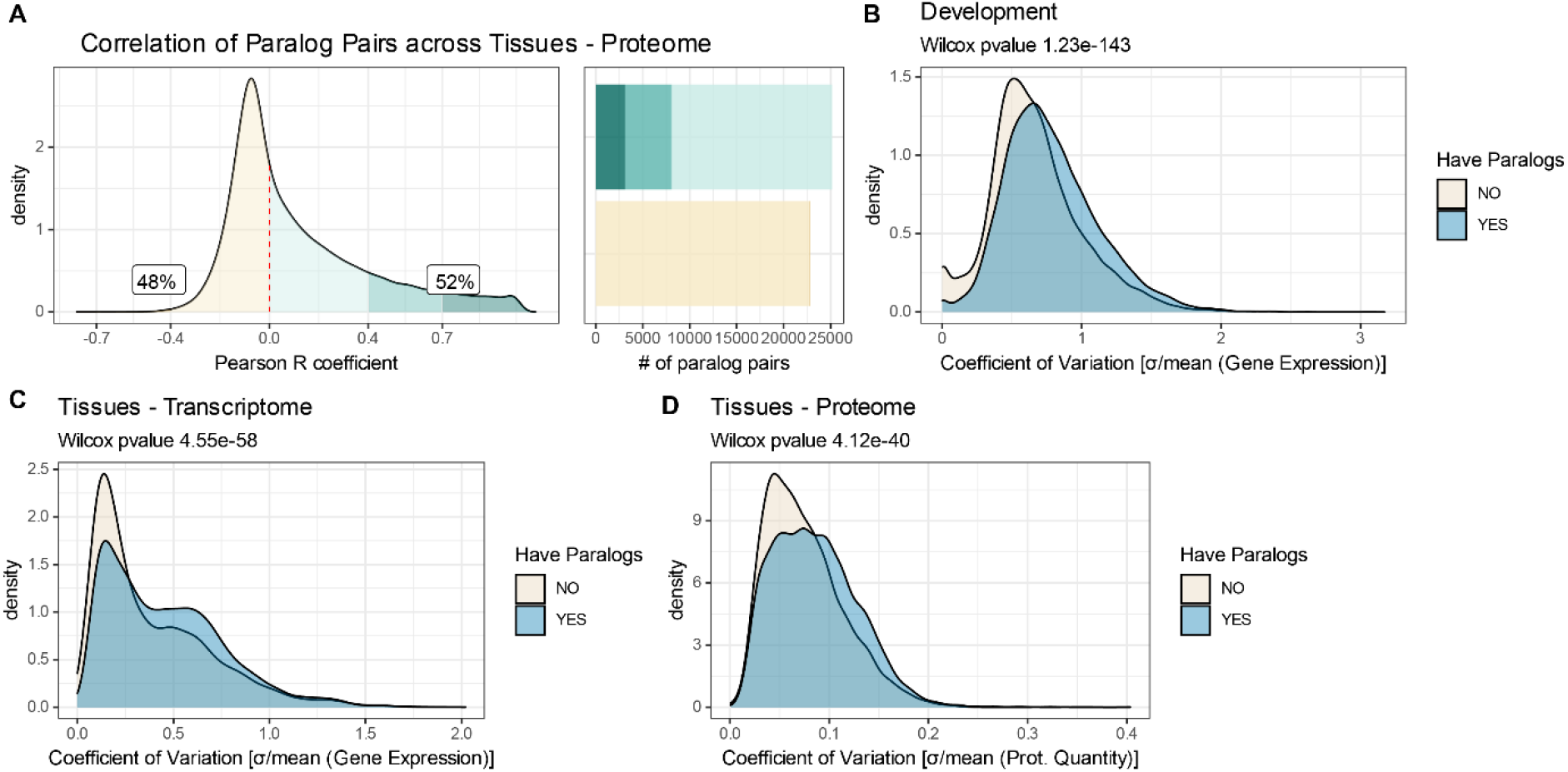
Paralogs contribute to transcriptome and proteome variability during development and across tissues. A - Density distribution of paralog protein pairs Pearson correlation across human tissues. Colored areas highlight specific correlation intervals. Labels indicate the percentage of paralogs that are positively correlated (R>0) and negatively correlated (R⇐0). Barplots indicate the numbers of paralog pairs present in each category. B-D - Coefficient of variation of genes that have paralogs (blue) and genes that do not have any (grey). Density distributions are shown for Zebrafish development (B), transcript across tissues (C) and protein across tissues (D).

**Supp. Fig2.**
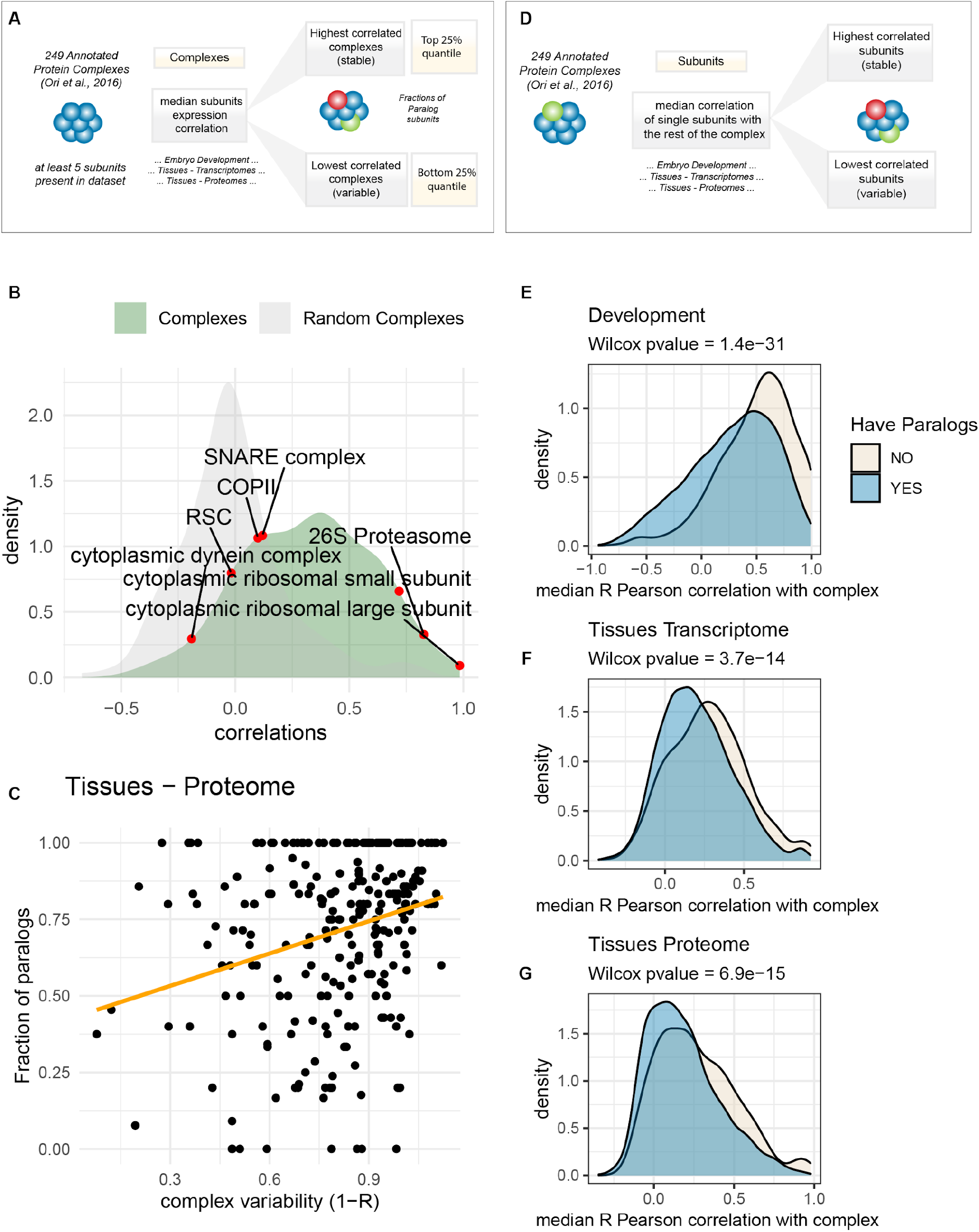
Paralogs contribute to the variability of protein complexes. A - Scheme of the protein complex analysis workflow. B - Density distributions of median Pearson correlation coefficient of the zebrafish development data for protein complexes (green) and a randomized set of protein complexes (grey) obtained by randomly assigning genes to protein complexes of the same size as the original set. Selected complexes displaying high or low correlations are highlighted on the distribution. C - Relationship between paralogs content (fraction of subunits that have paralogs in the genome) and complex variability. Complex variability is expressed as 1-R, where R is the median Pearson correlation of expression between all complex subunits. Based on proteome data from human tissues. D - Scheme of the Analysis - For each subunit of the 249 protein complexes in each dataset, we calculated the median Pearson correlation coefficient with all the other subunits of the same complex. The median Pearson correlation coefficient is used to assess whether each subunit is stoichiometrically stable (high correlations) or variable (low correlations). E-G - Distributions of median Pearson correlation coefficient for single subunits against their complex. Subunits with paralogs (blue) show lower median correlation with their complex compared to subunits that do not have paralogs (grey). Results are shown for zebrafish embryo development (E), transcriptome (F) and proteome (G) across tissues.

**Supp. Fig3.**
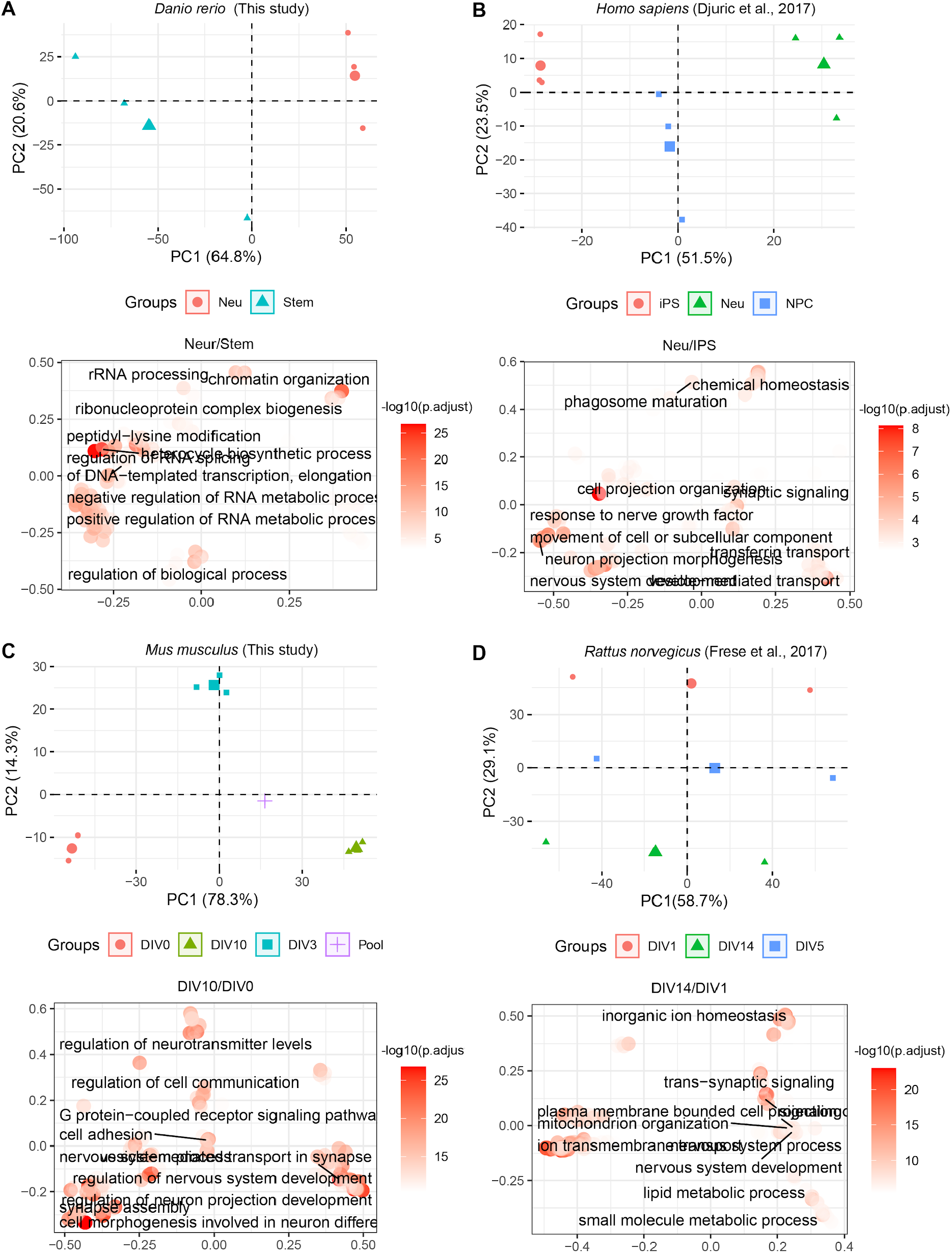
Proteome data of neuronal differentiation. A-D - Principal Component Analysis (PCA) of the proteomics data used in this study. Small symbols indicate the different replicates, large symbols indicate the centroid of each condition. Color and symbols indicate the different conditions considered. Below each PCA plot, the over representation analysis for “Biological Process” GO terms enriched among upregulated proteins (Log2 fold change >=0.58) against the rest of the quantified proteins is shown. In each plot, the x and y axis indicate semantic space separating the different terms, the color scale indicates the −log10(adjusted p value) of the Fisher test. DIV= differentiation *in vitro* day, iPSC=induced pluripotent stem cell, Neu= Neurons, NPC= neuronal precursor cell, Stem= undifferentiated stem cell.

**Supp.Fig4.**
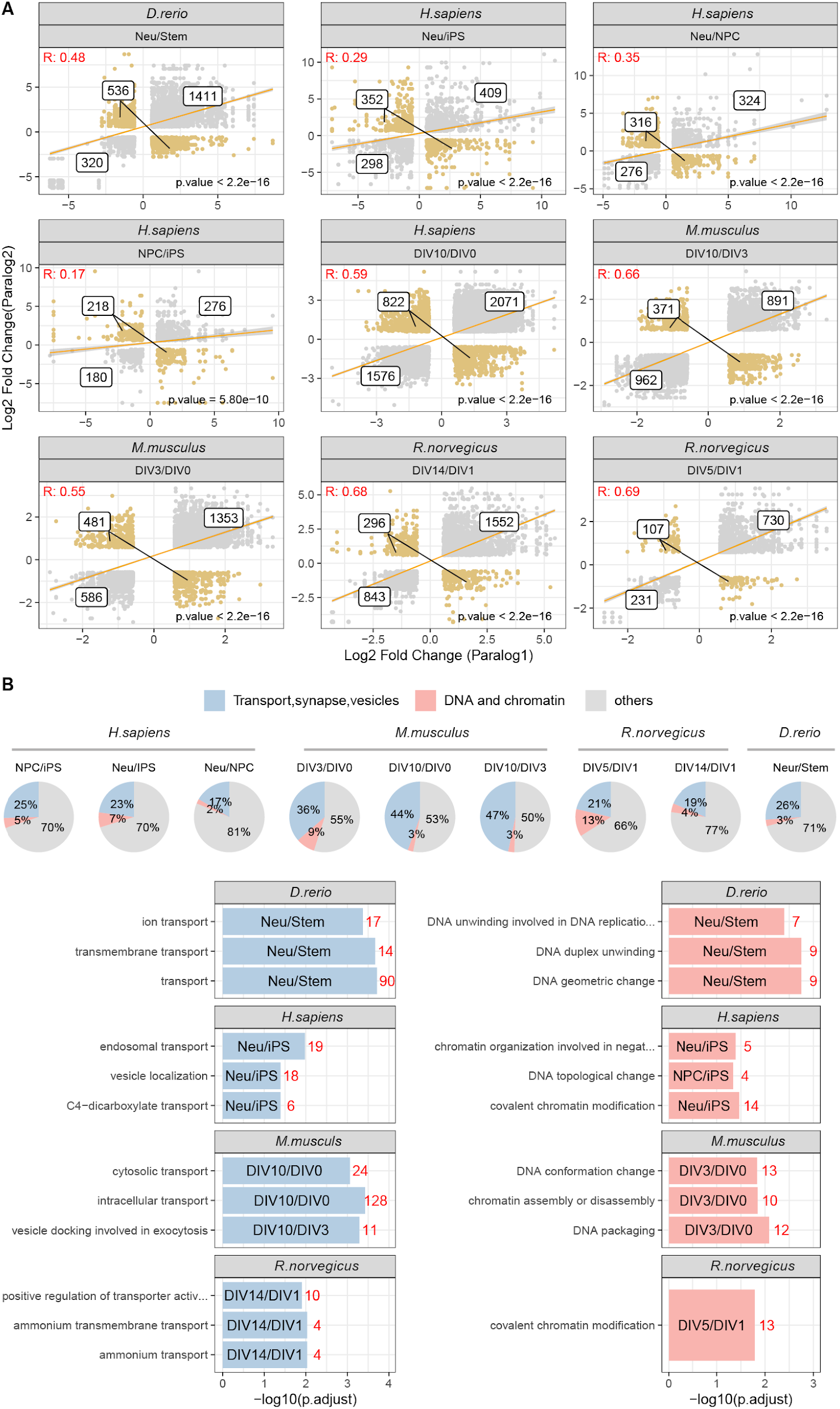
Co-regulation of paralog pairs during neuronal differentiation. A - Scatter plots compare Log2 Fold Change of paralog pairs during neuronal differentiation. Co-regulated paralog protein pairs are shown as grey dots, while paralog pairs regulated in opposite directions are shown in orange. A linear regression line is shown in yellow. Only differentially abundant proteins are shown for each dataset. Labels represent the number of unique paralog pairs present in each of the quadrants. B - GO terms enriched among paralog proteins that show opposite regulation during neuronal differentiation (indicated in orange in panel A) against the background of all differentially expressed paralog pairs. The pie plots indicate the percentage of differentially regulated paralog annotated with GO terms related to transport, vesicle, and synapses (in blue) and proteins annotated with GO terms related to DNA and chromatin in red. The barplots indicate GO Enrichment (ORA) for the specific GOTerms highlighted in the different pie plots. x-axis indicated the −log10(adjusted p value) for the enrichment. The red numbers indicate the amount of significant proteins present in each category.

**Supp.Fig 5.**
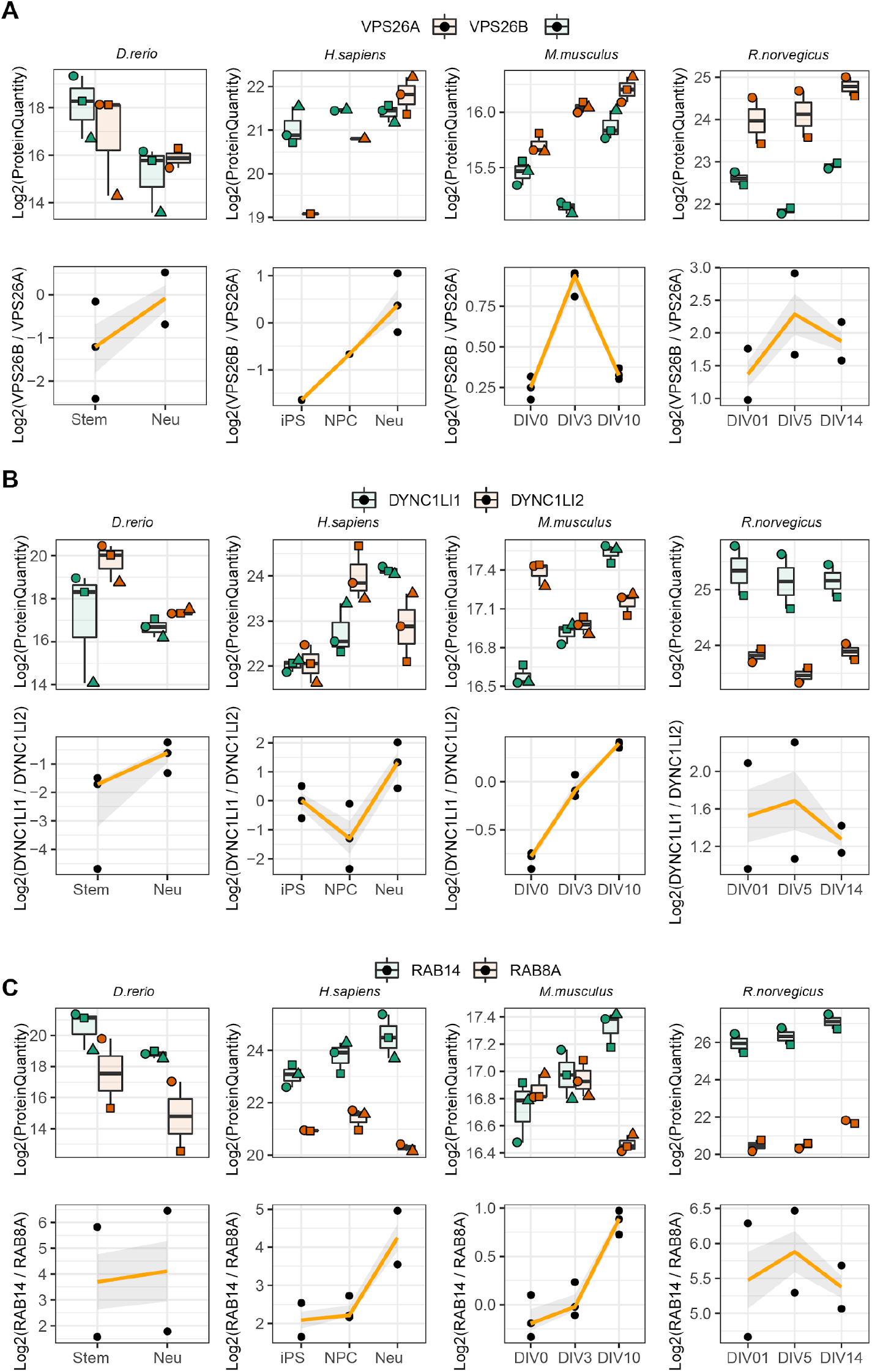
Changes of paralog ratios during neuronal differentiation. A-C - Protein abundance profiles for selected pairs of paralogs across datasets. Boxplots indicate Log2 protein quantities, across different replicates, while line plots (bottom) indicate the ratios between the two paralogs. In the top panel, shapes indicate paired replicate experiments. In the bottom panel, orange lines indicate the mean paralog ratio across replicates, and the shaded area represents 50% confidence intervals.

**Fig Supp 6.**
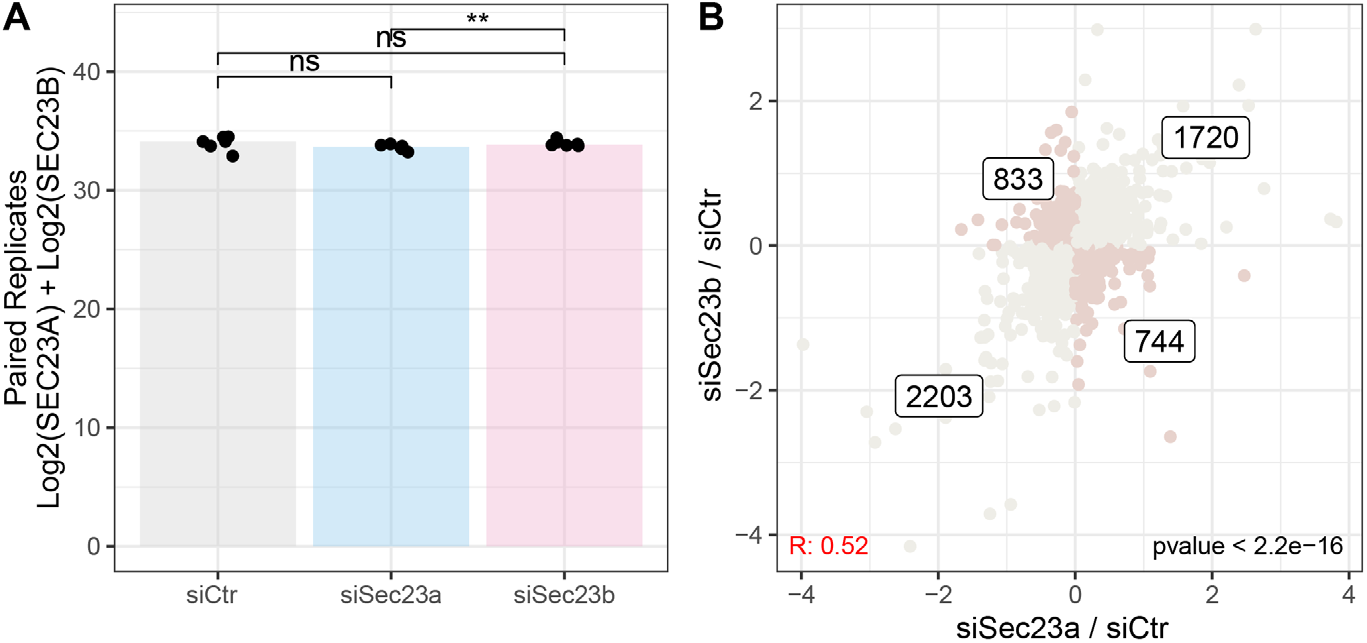
Effects of SEC23A/B knockdowns. A - Barplot shows the summed Log2 protein quantities of SEC23A and SEC23B following different siRNA treatment. Asterisks indicate significance for the paired two-sample Wilcoxon tests between conditions. ** p⇐0.01, ns=not significant. n=6. B - Scatterplot showing the relationship between the proteome changes induced by siSec23a (x-axis) and siSec23b (y-axis) against the siCtrl. Proteins affected in a similar way by the knockdown of the paralogs are shown in lightgrey, while the darker dots highlight proteins that are affected in opposite directions. The number of proteins present in each quadrant is indicated.

